# Bioframe: Operations on Genomic Intervals in Pandas Dataframes

**DOI:** 10.1101/2022.02.16.480748

**Authors:** Open2C, Nezar Abdennur, Geoffrey Fudenberg, Ilya Flyamer, Aleksandra A. Galitsyna, Anton Goloborodko, Maxim Imakaev, Sergey V. Venev

## Abstract

**Motivation:** Genomic intervals are one of the most prevalent data structures in computational genome biology, and used to represent features ranging from genes, to DNA binding sites, to disease variants. Operations on genomic intervals provide a language for asking questions about relationships between features. While there are excellent interval arithmetic tools for the command line, they are not smoothly integrated into Python, one of the most popular general-purpose computational and visualization environments.

**Results:** *Bioframe* is a library to enable flexible and performant operations on genomic interval dataframes in Python. *Bioframe* extends the Python data science stack to use cases for computational genome biology by building directly on top of two of the most commonly-used Python libraries, *numpy* and *pandas*. The *bioframe* API enables flexible name and column orders, and decouples operations from data formats to avoid unnecessary conversions, a common scourge for bioinformaticians. Bioframe achieves these goals while maintaining high performance and a rich set of features.

**Availability and implementation:** *Bioframe* is open-source under MIT license, cross-platform, and can be installed from the Python package index. The source code is maintained by Open2C on Github at https://github.com/open2c/bioframe.

## Introduction

Genomic interval operations are fundamental to bioinformatic analyses. These operations can be used to answer questions that include: Where is the closest enhancer to a gene of interest? How do chromatin states change across cell types? Which repeat elements contain binding sites for transcription factor motifs? Which annotations are enriched for SNVs associated with various diseases? Which promoters contain eQTLs? Given the ubiquity of these sorts of queries in genomic analysis, specialized interval arithmetic tools have been developed for the command line (Quinlan and Hall 2010; Neph et al. 2012), which operate on genomic interval text files such as the BED format. To facilitate interactive and programmatic use cases, there are also implementations for popular programming environments, including Python (Stovner and Sætrom 2020; Dale, Pedersen, and Quinlan 2011; Russell and Fiddes 2021), and R (Lawrence et al. 2013; Lee, Cook, and Lawrence 2019; Akalin et al. 2015).

The rich and robust set of data science and machine learning libraries in Python make it a popular choice for computational biology and data science more broadly. Core libraries in the Python data science stack, including *pandas* (Reback et al. 2020), *numpy* (Harris et al. 2020), *matplotlib* (Hunter 2007), and *jupyter* notebooks (Kluyver et al. 2016) offer nearly seamless integration. Other disciplines have developed tools to leverage the rich python data science infrastructure, e.g. GeoPandas (Jordahl 2014), SpatialPandas (Pothina, Pevey, and Lewis 2020), BioPandas (Raschka 2017) (for molecular structures). However, Python libraries for genomic intervals do not yet meet this high standard of integration.

Current Python packages providing support for genomic interval operations have limitations that impede smooth integration into Python data science workflows. For example, *pybedtools (Dale, Pedersen, and Quinlan 2011)*, the wrapper for *bedtools (Quinlan and Hall 2010)*, relies on interconversion between in-memory objects and text files stored on disk because data processing is delegated to a command line program. Furthermore, it inherits an API designed for the command line, and is restricted by rigid genomic interval schemas designed for storage (e.g. Browser Extensible Data (BED) files^1^). This leads to lower expressivity and less flexibility than what can be accomplished using Python-native (e.g. *numpy* and *pandas*) data structures and operations, as well as terse arguments with unintuitive names (e.g. -wao). The consequences include decreased performance, more boilerplate code, loss of metadata (such as column names), and code that is more difficult to read and debug. More recently, *PyRanges* (Stovner and Sætrom 2020), addresses many of these shortcomings, in particular by providing a 10-50x speed increase, but still has an API that is somewhat insulated from the data science stack. Conversions are required to switch between the custom *PyRanges* object used to perform genomic interval operations and standard *pandas* DataFrames, and *PyRanges* columns have relatively strict naming conventions.

The growth of the Python data science ecosystem presents an opportunity to re-imagine the implementation of genomic interval operations to smoothly interface with the full data science stack. Integration with this ecosystem can enable new avenues for data visualization, modeling, and insight into genomic data. Here we present ***bioframe*** (https://github.com/open2c/bioframe), a Python library for operating on genomic interval sets built directly on top of the *pandas* data analysis library. As a result, *bioframe* is fast and Pythonic, providing immediate access to a rich set of DataFrame operations, enabling complex workflows as well as rapid iteration, inspection and visualization of genomic analyses.

### Design Principles

The goal of *bioframe* is to enable in-memory, programmatic workflows on sets of genomic intervals using *pandas* DataFrames, integrating smoothly with the Python data science stack. With this in mind, we aimed to:

1. Reuse existing Python data structures. We encode interval sets using *pandas* DataFrames and avoid introducing new custom objects, e.g. ones based on interval trees.
2. Reuse existing Python methods. We delegate generic DataFrame operations to *pandas*, whenever possible, and aim at the principle of least_surprise^2^ for experienced *pandas* and *numpy* users.
3. Permit flexible data schemas. We avoid hard-coded column names, numbers and orderings.

### Definitions

To implement genomic interval operations in Python, we required formal definitions, which we did not readily find in one place in the literature. We thus put together definitions, starting from an interval (https://bioframe.readthedocs.io/en/latest/guide-definitions.html). We implement these definitions as specifications (https://bioframe.readthedocs.io/en/latest/api-validation.html) for properties of genomic interval dataframes.

While aligning reads to the full set of scaffolds in an assembly is typically advisable^3^, using a subset of scaffolds and/or breaking scaffolds into semantic subintervals (e.g. chromosome arms) is often crucial for downstream genomic analyses. In *bioframe* we thus introduce the concept of a ***genomic view*** to specify a unique genomic coordinate sub-system. A genomic view is an ordered set of uniquely-named non-overlapping genomic intervals known as *regions*. There can be more than one region from the same scaffold and multiple scaffolds represented in a view. Defining a view allows a user to focus analysis on a well-characterized portion of an assembly and specify the order of scaffolds and regions for effective visualization. Indeed, defining a view for downstream analysis can be more important for non-model organisms, where assembly quality is often lower and thus requires judicious choice of order and subset of scaffolds to analyze.

### Implementation

*Bioframe* is implemented using the machinery of *numpy* and *pandas*, making for a lightweight set of dependencies and avoiding the need to compile custom C extensions or other external libraries. For example, to determine overlaps between intervals (*bioframe.overlap*), as well as find pairs of nearby intervals (*bioframe.closest*), *bioframe* uses sorting-based algorithms. First, intervals are split into subsets by chromosome and optional columns like ‘strand’ (*pandas.groupby*). These interval subsets are then sorted (*numpy.lexsort*), and overlaps (or neighbors) are detected via bisection search operations (*numpy.searchsorted*). All user-facing operations are imported into the base bioframe namespace.

Building directly on *pandas* allows *bioframe* to readily generalize the genomic interval model used for Browser Extensible Data (BED) files^4^. *Bioframe* requires only genomic coordinate columns -- the equivalent of ‘chrom’, ‘start’, ‘stop’-- with flexible names, for a valid BED-like DataFrame, or ‘bedframe’. Almost any number of additional annotations can be added to a set of intervals.

### Functionality

The core genomic interval operations in *bioframe* are: *overlap, cluster, closest, and complement. bioframe* additionally provides frequently-used operations that can be expressed as combinations of these core operations and *pandas* dataframe operations, including: *coverage, count_overlaps, expand, merge, select, subtract, setdiff, trim*. Building from the definition of genomic views, *bioframe* provides functions to: assign intervals in a bedframe to regions in a genomic view, *assign_view*, and sort a bedframe based on the order of regions specified in a view, *sort_bedframe*.

Building on *pandas* enables flexible control over column usage and selection. *Bioframe* includes a context manager for setting default column-names for genomic coordinate columns. This flexibility shines when dealing with BED-like files that can have variable headers or conventions for the genomic coordinate columns (e.g. ‘chrom’ or ‘chromStart’ or ‘chr’ or ‘CHR#’), and variable orders of other interval metadata columns (e.g. ‘score’, ‘color’, or ‘strand’). Since operations like overlap are performed following a *pandas* groupby, *bioframe* also flexibly generalizes genomic operations that consider strand to any list of common columns present in a pair of dataframes.

In addition to these features, *bioframe* provides functions for dataframe construction, checks, string operations, and I/O. For example, there are wrappers and schemas for reading and writing common binary and text genomic file formats to and from dataframes.

### Performance

We profiled speed and memory usage for typical use cases using *bioframe*, and compared performance with that of *pybedtools* and *PyRanges*. We intersected sets of random genomic intervals stored as *pandas* DataFrames, accounting for format conversions needed in *pybedtools* and *PyRanges*. For overlaps of up to 3*10^6^ intervals, *bioframe* and *PyRanges* have comparable speeds (**Figure 1A**), while pybedtools can be more than 100x slower. The memory consumption of *bioframe* is almost exactly 2x higher compared to *PyRanges* and *pybedtools* (**Figure 1B**) because it stores coordinates as 64-bit integers, motivated by the emergence of sequenced genomes with >1e9 base pairs per chromosome (e.g. axolotl (Nowoshilow et al. 2018), lungfish (Meyer et al. 2021)). We note that for chained operations, *PyRanges* offers further speedups by caching per-chromosome interval tables, with the tradeoff of storing intervals as a custom object with its own API layer. To conclude, both libraries offer reasonable performance (<1s for 10^5^ intervals). For much larger sets of genomic intervals (>10^7^), users may also want to consider other high-performance options (Li and Rong 2021; Neph et al. 2012).

**Figure 1.**
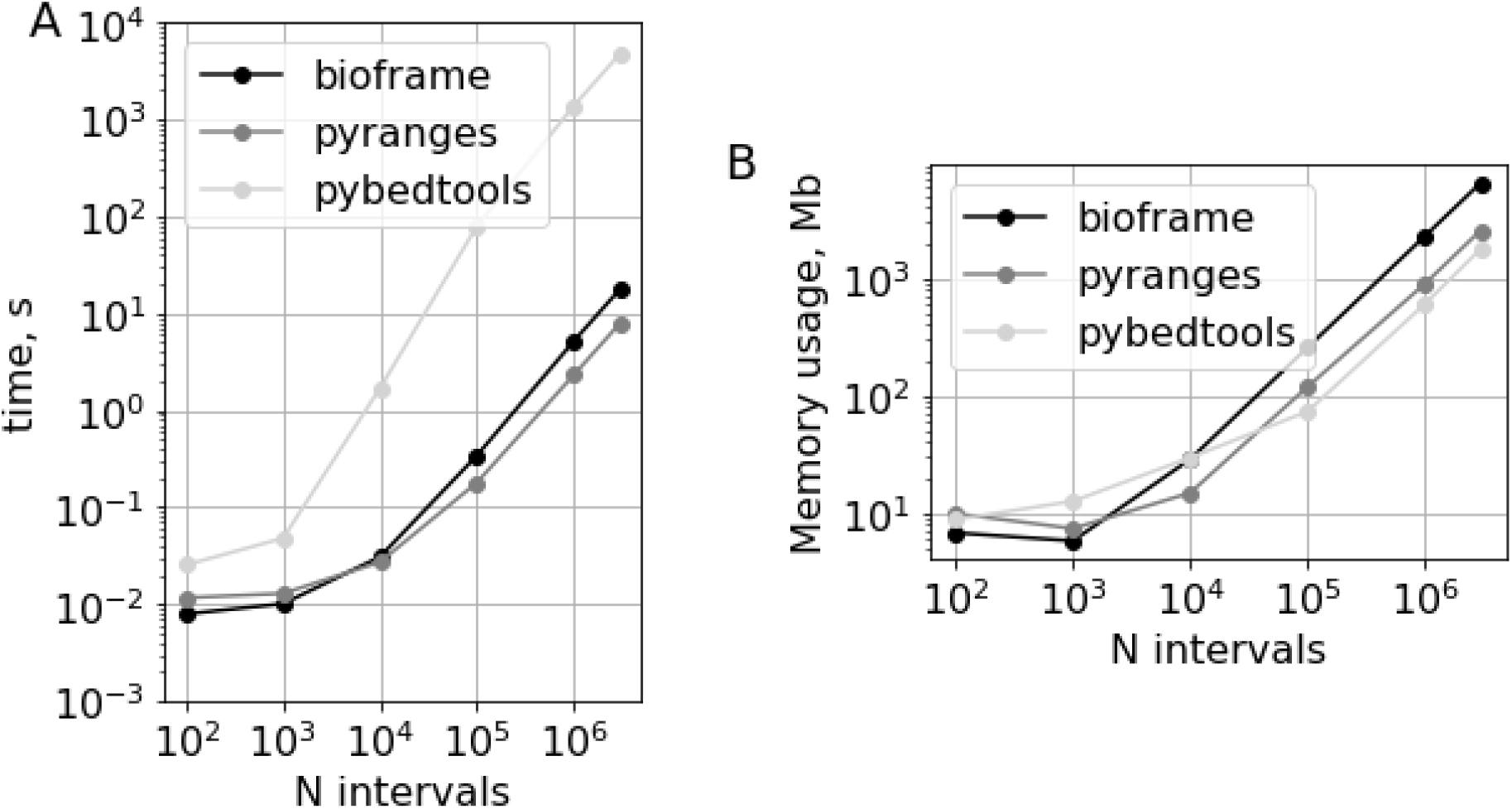
Performance comparison of *bioframe, PyRanges*, and *pybedtools* for detecting overlapping intervals between pairs of dataframes of randomly generated genomic intervals. (A) Run time and (B) peak memory consumption of *bioframe **overlap*** vs. *PyRanges **join*** show comparable performance up to millions of intervals. *Pybedtools **intersect*** shows slower performance and comparable memory usage. Code for this performance comparison is available at https://bioframe.readthedocs.io/en/latest/guide-performance.html

## Discussion

In summary, *bioframe* provides a pure Python library for genomic interval operations. *Bioframe* presents a Python-centered API for these operations, as opposed to inheriting syntax from the command line. Working in Python with pandas dataframes enables flexible generalization of the BED format, including flexible naming for genomic interval columns. *Bioframe* has already proven useful for pandas-heavy genomic workflows, like *cooltools* (Venev et al. 2021). In the future, providing tools specific to binned genomic intervals, paired genomic intervals, and out-of-core dataframe operations (e.g. with Dask (Rocklin 2015) or Modin (Petersohn et al. 2020)) would be valuable extensions to *bioframe*.

## Acknowledgements

The authors thank Sameer Abraham, Luis Chumpitaz, George Spracklin, and Aafke van den Berg for repository suggestions and Endre Bakken Stovner for helpful comments. AG is supported by IMBA and the Austrian Academy of Sciences (OeAW). GF is supported by R35 GM143116-01. IF acknowledges funding support from the MRC University Unit grant MC_UU_00007/2.

## Author Contributions

We welcome issues and questions on github https://github.com/open2c/bioframe/. For questions about array operations / implementation, tag AG @golobor, for questions about front-end interval operations, tag GF @gfudenberg, for questions about file I/O and resources tag NA @nvictus. IF, AAG, MI, and SV made contributions as detailed in the Open2C authorship policy guide. All authors are listed alphabetically, read, and approved the manuscript.

## Footnotes

1 https://genome.ucsc.edu/FAQ/FAQformat.html#format1

2 https://en.wikipedia.org/wiki/Principle_of_least_astonishment

3 https://lh3.github.io/2021/04/17/concepts-in-phased-assemblies

4 https://samtools.github.io/hts-specs/BEDv1.pdf

## Notes

https://github.com/open2c/bioframe

### Competing Interest Statement

The authors have declared no competing interest.

